# Pre-existing Technological Core and Roots for the CRISPR Breakthrough

**DOI:** 10.1101/329706

**Authors:** Christopher L. Magee, Patrick W. Kleyn, Brendan M. monks, Ulrich Betz, Subarna Basnet

## Abstract

This paper applies objective methods to explore the technological origins of the widely acclaimed CRISPR breakthrough in the technological domain of genome engineering. Previously developed patent search techniques are first used to recover a set of patents that well-represent the genome editing domain before CRISPR. Main paths are then determined from the citation network associated with this patent set allowing identification of the three major knowledge trajectories. The most significant of these trajectories for CRISPR involves the core of genome editing with less significant trajectories involving cloning and endonuclease specific developments. The major patents on the core trajectory are consistent with qualitative expert knowledge of the topical area. A second set of patents that we call the CRISPR roots are obtained by finding the patents directly cited by the recent CRISPR patents along with patents cited by that set of patents. We find that the CRISPR roots contain 8 key patents from the genome engineering main path associated with restriction endonucleases and the expected strong connection of CRISPR to prior genome editing technology such as Zn finger nucleases. Nonetheless, analysis of the full CRISPR roots shows that a very wide array of technological knowledge beyond genome engineering has contributed to achieving the CRISPR breakthrough. Such breadth in origins is not surprising since “spillover” is generally perceived as important and previous qualitative studies of CRISPR have shown not only technological breadth in origins but scientific breadth as well. In addition, we find that the estimated rate of functional performance improvement of the CRISPR roots set is about 9% per year compared to the genome engineering set (˜4 % per year). These estimates indicate below average rates of improvement and may indicate that CRISPR (and perhaps yet undiscovered) genome engineering developments could evolve in effectiveness over an upcoming long rather than short time period.

## Introduction

Genome engineering has been one of the promising biomedical approaches studied in the past few decades. Just 5 years ago, CRISPR-Cas9 emerged as a much more economical, practical and generalizable genome editing technology. Since then it has become popular to refer to CRISPR as the most important biotechnology breakthrough of the 21^st^ century (1) and as one of the two (PCR being the other) most important biological technologies of the past 50 years (2). Genome engineering is genetic engineering in which DNA is inserted, deleted, modified or replaced in the genome of a living cell or organism. Since there is not a consensus about differentiation, we -and most others- use genome editing as a synonym for genome engineering. There is consensus that CRISPR-an acronym for Clustered Regularly Interspaced Short Palindromic Repeats -and Cas9 (CRISPR-associated protein 9) is the nomenclature for the signature protein for type II CRISPR systems that, directed by guide RNAs, cleaves DNA in a sequence-dependent manner. CRISPR (and Cas9) were discovered in bacteria (3-7) where they form the backbone for very effective viral resistance systems in numerous species setting the stage for other uses (8). Lander in a paper retracing the history (9), Doudna and Sternberg in a memoir and historical book (10) and more recently Urnov (11) all do an excellent job of covering the *many strands of globally-dispersed scientific work* (including discovery of CRISPR, its role as an adaptive immune system, experiments confirming the CRISPR role and showing use of a nuclease, adapting findings from earlier genome editing techniques, sorting out the importance of the various Cas proteins especially Cas9, cRNAs- or CRISPR RNA complexes, discovery of tracRNA, reconstituting CRISPR in a distant organism, studying CRISPR in vitro) *essential* to the initial sets of CRISPR patents. It is particularly interesting that many of these scientific research studies were undertaken for reasons having no biomedical intention (and often not focused on CRISPR or genome editing). This scientific story is fundamental to the emergence of CRISPR and the Lander article, the Doudna and Sternberg book and the Urnov article are recommended if one wants to understand it (9, 10, 11). This paper does not emphasize the scientific literature but instead focuses on the patent literature associated with genome editing and CRISPR. We note that patents do cite scientific papers but scientific papers almost never cite patents so study of patents is an important element in the emergence and development of any technology. We also note that there are several legal conflicts about patents in this area and that the growth of relevant patent applications has “exploded” since 2012. The rapid growth and the legal conflicts do not –in our judgement-eliminate the usefulness of assessing the technological core and roots of CRISPR in the patent system.

There is extensive development of methods, based upon analysis of patents that are aimed at improving understanding of technological developments such as CRISPR. This paper (to our knowledge the initial attempt to analyze CRISPR in this way) will utilize two promising analytical frameworks-the first is usually called main path (or knowledge trajectory) analysis and the second is called rate of improvement estimation.

Main path analysis began with Hummon and Doreian’s technique for analysis of citation networks of scientific papers and their initial application was to the development of DNA theory from 1820 to 1965 (12). They developed the methodology and demonstrated it by identifying the key papers in this knowledge trajectory. Verspagen (13) and Mina et al (14) then adapted main path analysis for technological knowledge trajectories by applying the Hummon and Doreian technique to the patent citation network for fuel cells (13) and coronary artery disease treatment (14). The technique has been extended (15) and applied to several other technological domains (16, 17) including telecommunication switching, solar photovoltaics, desalination and others. A technique for obtaining relevant and relatively complete patent sets for characterizing domains developed by Benson and Magee (18, 19) proved useful in main path analysis (17) and is the starting point for gathering patents in the present work.

Empirical study of the change in technological performance with time (20-30) has shown that the exponential dependence first noted by Gordon Moore (20) applies (with ample noise) to all domains studied. It is also clear that the exponent (or % change per year) varies among technological domains from ˜1.5% per year to ˜65% per year (28, 30). Obtaining empirical estimates for any given domain is problematic and at best extremely time consuming but recent work (31-34) has resulted in reliable estimates based upon representative sets of patents for the domain of interest. Indeed, Triulzi et al (33) have shown that the most reliable estimate of performance improvement rate is based upon analysis of the same patent citation network used to determine knowledge trajectories. Domains that improve more rapidly carry more than their share of the total information flow on the overall patent citation network; that is, their patents have higher average information centrality.

The extremely high interest in and potential for CRISPR along with the patent analysis methods just mentioned led to the formulation of two research objectives guiding the current research. The first research objective involves determining what the patent record shows about the relationship of CRISPR to prior technology- particularly pre-existing genome engineering technology. The second research objective is to estimate the rate of improvement in performance of genome engineering and CRISPR.

### Collection of data

#### Genome engineering patent set

The current research utilizes two sets of US patents for the quantitative empirical study. The first set of patents represent the genome engineering domain and are retrieved using the Classification Overlap Method (COM) (18,19) which utilizes two different classification systems to obtain highly relevant patents. In this study, the COM procedure was implemented in 5 steps. (step 1) Preparation of Pre-set patents: This step can utilize representative key inventors, assignees, or patents. In the current study, we utilized 58 key patents found by searching for some known inventors of genome editing technologies. (step 2) Identification of classes in two distinct classification systems: we chose the US Patent Classification (UPC), and the Cooperative Patent Classification (CPC) as the systems. Mean Precision-Recall (18, 19) was used as a metric to identify the relevant classes in UPC and in CPC. (step 3) Patents that are common to classes in UPC and in CPC identified in Step 2 are retrieved; (step 4) Test of relevancy: A sample of retrieved patents (most cited 100 patents and 200 randomly selected patents from the remaining) were read (mostly just titles and abstracts) by the investigators to determine relevancy of the patent set. (step 5) For completeness, the classes were checked to ensure that more than 75% of the 58 key patents were included in the retrieved set of patents.

To generate the final genome engineering patent set, the steps above were applied to all granted US patents from 1970/01/01 to 2018/01/15 available in Patsnap, a commercial patent database (35). The 58 key patents for Step 1 were identified by a domain expert through literature review of patents found by searching for known major participants in genome editing technologies. The 58 patents uncovered include 28 patents related to zinc finger nuclease (ZFN), 8 patents for transcription activator-like nucleases (TALEN), 6 patents for meganuclease and 16 patents for CRISPR. An in-depth study of a sample of patents in the genome editing patents showed that significant number of the patents were classified in many classes. For example, patent number US8865406 is classified into 14 UPC classes, which is unlike what is typically seen in other technological domains such as Solar Power, Batteries, and Integrated Circuits (average is 3.2 UPC classes). Further, we also observed that the Mean-Precision Recall value of UPC and CPC classes decayed slowly as compared to other domains. This implied that potentially relevant patents were widely dispersed across many classes both in UPC and in CPC. This made it necessary to include multiple classes both in UPC and in CPC to attain adequate coverage of patents and dictated that reading titles and abstracts was done in multiple iterations.

Fig. 1 shows the classes considered to retrieve genome engineering patents, which are decomposed into three components for readability: The first component consists of patents related to ZFN, TALEN, and meganuclease. As shown in Figure 1, this component uses four classes from UPC and four from CPC. The second and third components consist of patents related to CRISPR, and uses a Ribonuclease class both in UPC and in CPC. We note here that COM utilizes two classification systems to identify patents in a domain, as the co-occurrence in two classes in different systems leads to highly relevant patents (18, 19). Since mid-2015, the USPTO has stopped classifying US patents using UPC classes. Therefore, we split the period into prior to mid-2015, and after mid-2015, so we may still gain the advantage of COM’s effectiveness in yielding highly relevant set of patents for the period before mid-2015. The third component utilizes only the CPC class. Using the classes and the time period considered (1970/1/1-2018/1/15), we retrieved 1373 patents. Hereafter, this group of patents is referred to as the *genome engineering patent set*. The set covers 78% of 58 patents in the Pre-set patents. Out of 28 Zn finger patents, 18 were recovered; for Talen 6 out of 8; for Meganuclease 5 out of 6; and for CRISPR 16 out of 16 (See Fig 1C).

**Fig. 1.**
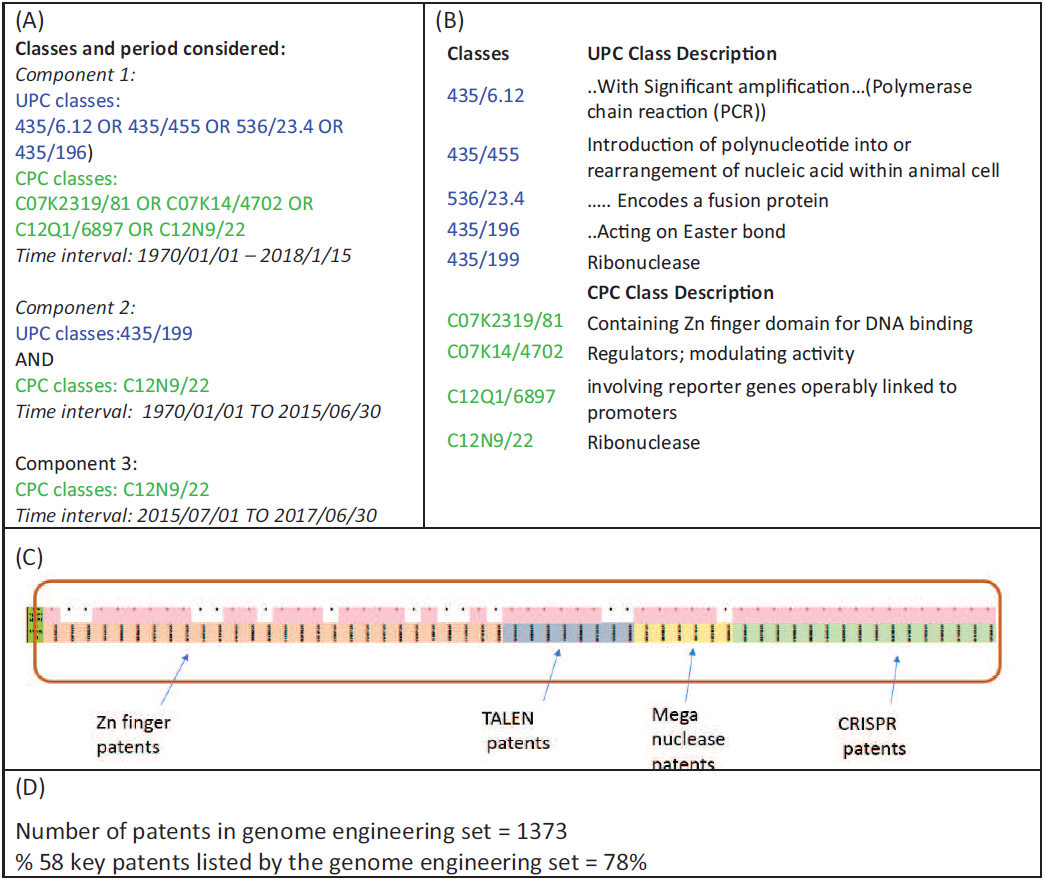
Application of Classification of Overlap Method (COM). (A) UPC and CPC classes and time period used to implement COM; (B) Description of UPC and CPC classes; (C) Visual depiction of the 58 patents in the pre-set in the classes selected, an indication of completeness. White spaces indicate the patents not retrieved in this set; (D) Total patents retrieved and percentage of 58 key patents covered by the Pre-CRISPR patents.

Patenting activity for genome engineering occurred at a steady pace from 1999 until 2012 with about 40-60 patents granted per year (see Figure 2A). The patenting activity, however, greatly accelerated recently, doubling to about 115-120 patents for 2016 and 2017 with the accelerated pace due to pursuit of CRISPR technology. Figure 2B shows the top 10 assignees for the genome engineering patent set.

**Fig 2.**
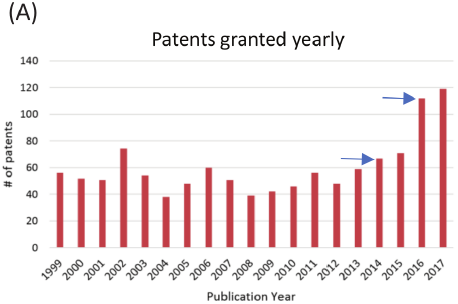
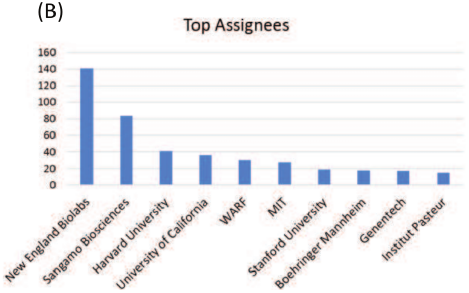
Patenting activity for genome engineering patent set. (A) patents granted yearly 1999-2017; (B) Top 10 assignees (with formal names) New England Biolabs, Sangamo Biosciences, Harvard University (President and Fellows of Harvard College), University of California (The Regents of The University of California), WARF (Wisconsin Alumni Research Foundation), MIT (Massachusetts Institute of Technology), Stanford University (The Board of Trustees of The Leland Stanford Junior University), Boehringer Mannheim (Boehringer

#### CRISPR Roots Patent Set

This study also undertook a direct generational study of the citation network emanating from the CRISPR patents. The creation of a new CPC patent class by the USPTO during 2017 - specifically to contain CRISPR patents- defined a useful starting point to find current CRISPR patents. As of January 14, 2018, this CPC patent class (C12N2310/20) contained 37 patents (granted between 1976/1/1 – 2018/1/15) which we call Generation 0 (in short Gen0 patents). We then retrieved the 112 granted patents cited by Gen0 patents (generation 1, in short Gen1). These 112 patents are those remaining after those cited that were already in Gen0 were removed, thus making Gen1 mutually exclusive. We then retrieved 1230 patents cited by Gen1 patents, but not belonging to Gen0 or Gen1, as Generation2 (in short, Gen2) patents. It is noted that there was no restriction as to what classes the cited patents in Gen1 and Gen2 belonged. These three subsets, Gen0, Gen1 and Gen2, in total 1379 patents, make up the patent network directly generated by citation cascade from CRISPR patents in 2 generations of citations. We designate this set of patents the CRISPR roots patent set and will use this terminology hereafter. Figure 3 shows descriptive information about the time dependence and ownership of this patent set.

**Fig 3.**
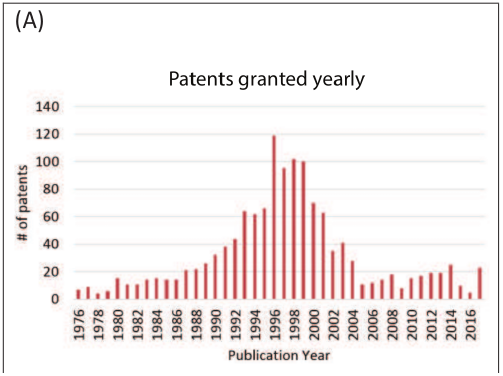
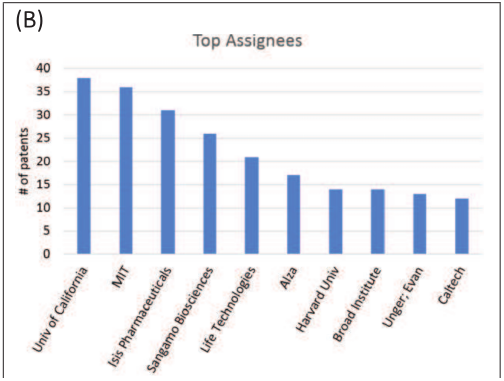
Patenting activity for CRIPSPR Roots set. (A) Patents granted yearly 1976-2017; (B) Top 10 assignees (with formal names): Univ of California (The Regents of The University of California), MIT (Massachusetts Institute of Technology), Isis Pharmaceuticals, Sangamo Biosciences, Life Technologies, Alza (Alza Corporation), Harvard Univ (President and Fellows of Harvard College), Broad Institute (The Broad Institute, Unger, Evan (Evan C. Unger), Caltech (California Institute of Technology).

Fig. 3A shows yearly patents granted from 1976 until 2017 for the CRISPR roots. Most of patents in the set were granted from the late 1980’s to the early 2000’s. This distribution over time is not surprising: about 89% of the patents in the set belong to Gen2 which represent the relatively older citations from Gen1. Fig 3B shows the top 10 assignees in the CRISPR roots.

#### Main path Methodology

The main path methodology provides the means to identify important patents in the technological domain and pathways through which the technological knowledge diffused in the domain. The method originated to understand the evolution of scientific fields through study of citations by scientific publications (12). The methodology was adapted and modified to investigate the evolution of knowledge in many technological domains (13-17). Most recently, the method has been optimized to produce simpler main paths, while capturing a greater number of important patents (17). Labeled as genetic backward-forward path (GBFP) analysis, the optimized method consists of four steps shown in Figure 4: assembling/collecting a patent set, constructing a citation network within the patent set, measuring knowledge persistence of the patents to identify genetically high-persistent patents, and tracing main paths (forward and backward) from the genetically high-persistent patents.

**Fig. 4.**
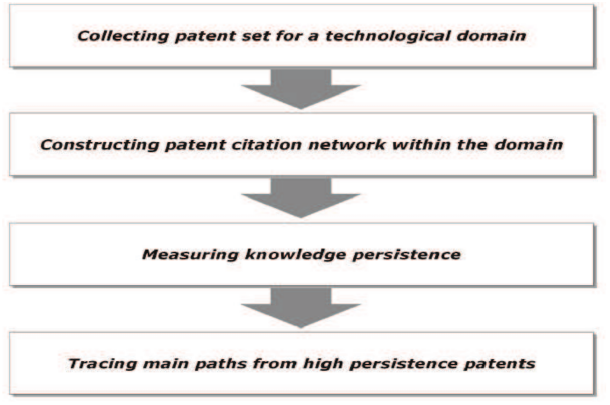
Steps for genetic backward-forward path analysis (GBFP) adapted from (17)

To implement the method for the genome engineering and CRISPR roots patent sets, the patent network is constructed using the citations made by the patents in the sets. It is noted that we consider citations only within the patent set; any citations outside the patent set are ignored. To estimate the persistence of knowledge (15,17) contained in each patent, the patent network is first ordered using the citations into *n* layers (visualize that the patents initially cited are on the left) and then knowledge persistence is estimated for each patent in the leftmost layer (layer 1). The process is repeated successively for the subsequent layers moving to the right (layers 2, 3, 4…) after eliminating all the layers to the left of the layer in question. This algorithm estimates two types of persistence values (0 to 1 after normalizing) for each patent in the network: global persistence (GP) and local persistence (LP). The GP of a patent is estimated to gauge the importance of a patent in the entire network whereas LP is estimated to gauge the importance of patents in each layer. The layer persistence plays a significant role in identifying and retaining important patents, which are recent, and hence, have not had a chance for their lineage to evolve. The high-persistent (GP > 0.3 and LP > 0.8) patents then become the origin for tracing for the main paths, both backward and forward (17). We adopt GBFP analysis to investigate the evolution of CRISPR within the genome engineering domain. Specifically, we use this methodology to identify important patents in genome engineering which preceded the CRISPR technology. By reading these important patents we are also able to identify technology clusters within genome engineering that preceded CRISPR.

#### Estimation of patent centrality and annual improvement rate (k)

The estimation of annual improvement rate for a set of patents starts- as does the main path method just described- with the patent citation network. The centrality of a patent is analogous to betweenness centrality in network analysis, and provides a measure of the influence a node, in our case the patent, has over flow of information (in our case, the technological knowledge) through the network. Our calculation of the information centrality can again be traced to Hummon and Doreian (12) and their introduction of search path node pairs (SPNP) as a metric to compute the centrality of a focal paper in a scientific paper citation network. The SPNP for a focal patent (say, patent B) in a patent citation network calculates the number of pathways originating from one patent (say, patent A) to another one (patent C) in the network *and* passing through the focal patent (patent B). The higher the number of pathways traversing through the focal patent the higher the centrality of the focal patent, indicating the importance of the focal patent in the patent citation network. Since each patent can be interpreted as containing some original technological knowledge, the centrality provides a sense of the importance of the original knowledge introduced by the focal patent for the downstream patents. Triulzi et al (33) normalized the SPNP to account for the variations inherent in the patenting system (for example, citation practices between fields, and particularly over time), which make raw centrality values of patents across domains and between two different time periods non-comparable. To control for these variations, the computed centrality of a patent is compared with the expected value of the centrality of the same patent in appropriately randomized models of the citation network (33). The centrality calculated was for the citation network of all US utility patents granted from 1976 until 2015. Triulzi et al further find that the mean normalized centrality of a patent set representing a specific technological domain is a reliable predictor of its annual rate of improvement (k). They arrive at this conclusion by a Monte Carlo cross-validation exercise between empirically observed k for the 30 diverse technological domains (28, 30) and their corresponding mean normalized centrality of the patent sets for the same 30 technological domains. Their regression model developed considering 30 technological domains is shown below:

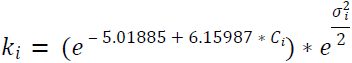

Where k_*i*_ represents the annual rate of improvement for domain *i*, C_*i*_ the mean normalized centrality of the patent sets for the domain *i*, and *σ* _*i*_ the standard deviation of C_*i*_. We have adopted their regression model to estimate the annual rate of improvement for the genome engineering and CRISPR roots patent sets. Indeed, we used the centrality calculations developed by Triulzi et al (33) for the patents in our patent sets to calculate the mean for the two sets which we treated as domains.

## Results

### Genome engineering main path

Figure 5 gives the results of applying the main path methods described in the previous section to the genome engineering patent set. The main path is a network with three principal components (GE1, 2 and 3). While all relate to the development of enzymes to bind and cleave DNA, GE1 and GE3 relate to the production of restriction endonucleases (REs) for general molecular biology applications whereas the larger GE2 path relates specifically to core genome editing development.

**Fig. 5.**
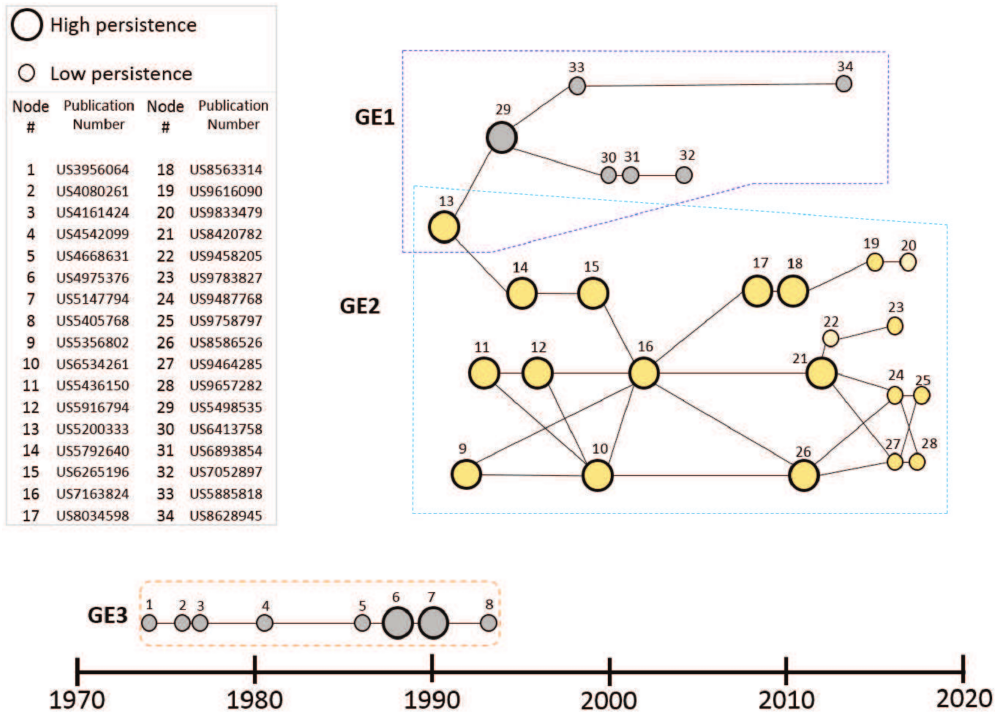
Main path results for genome engineering patent set. Three main paths (GE1, GE2 and GE3) have been identified. GE1: Cloning and restriction endonuclease (REs); GE2: core genome editing; GE3: Endonuclease and related enzymes. Labeled nodes represent patents and are identified in the side table with the patent number which allows one to search for and read the patent on various databases.

GE3 has the oldest patents dating to the mid-1970’s. The initial patents (1, 2 and 3), all assigned to Rikagaku, Japan, specify methods for purifying endogenous nucleases from bacterial cells. Subsequent patents in this path from the 80’s and the early 90’s relate to methods of producing specific REs.

Patent 13 (US5200333) belongs to GE1 and it also initiates GE2. This patent relates to improvements in methods of producing REs by selection of bacterial cells expressing methylase enzymes that confer resistance to the RE produced. The GE1 path extends this with further enhancements to the methodology of producing REs (patents 29, 30, 31,32) and applying these improvements for producing specific REs (patents 33 and 34). Most of the patents in GE1 are assigned to New England Biolabs indicating a significant role for them during the 1990’s improving the methods of RE production.

GE2 is the path of direct relevance to genome engineering. Based on the same improvements on RE production described in patent 13, GE2 combines these with major advances in creating synthetic novel REs that recognize rarer DNA targets using ZFNs and TALENs and ultimately CRISPR complexes that are applicable to genome engineering. This path is analyzed further in Figure 6 showing the key patents in the development of genome engineering that underlie the emergence of CRISPR.

**Fig. 6.**
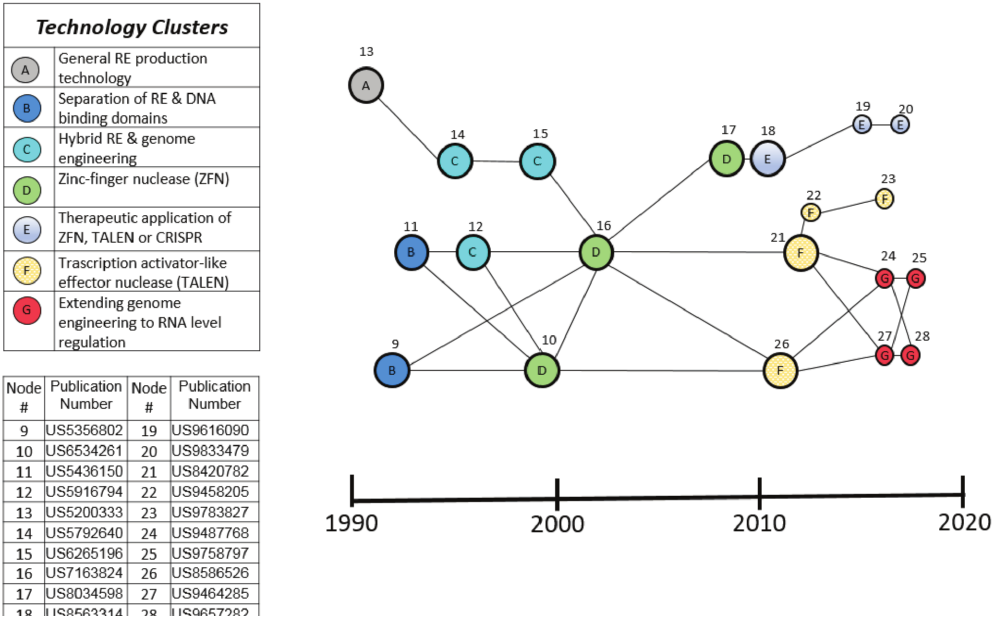
Technology clusters in GE2 main path. The patents in GE2 are identified in clusters of different technologies shown in the table in the upper left of the figure: (A) restriction endonuclease (RE) production technology; (B) separation of RE and DNA binding domains; (C) hybrid REs and genome engineering; (D) Zinc-finger nuclease (ZFN); (E) Therapeutic application of ZFN, TALEN, or CRISPR; (F) Transcription activator-like effector nuclease (TALEN); (G) Extending genome engineering to RNA level regulation. Nodes represent patents repeated from Figure 5 and the actual patent numbers are identified in the lower left legend in this figure.

Patents 9 and 11 (labeled cluster B in figure 6) from the early 90’s describe a fundamental step forward, taken by Chandrasegaran’s group at Johns Hopkins University, towards the goal of genome engineering: The separation of FokI restriction endonuclease (RE) into two distinct domains, one that binds its cognate target DNA sequence and the other containing the nuclease activity that cleaves DNA. This invention led to the possibility that the nuclease activity of FokI could be fused to alternative DNA binding domains to create so called “hybrid REs” with novel, and potentially rare DNA target sequences useful for genome engineering in large animal and plant genomes (36).

A significant challenge in producing hybrid REs in bacteria was that they were potentially lethal to their host bacteria if the latter contained target sequences in their genome (36). Patents 12 and 14 from the mid-90’s describe improvements to bacterial hybrid RE synthesis by co-expressing DNA ligases and/or expressing the hybrid REs on inducible plasmids to mitigate this risk. Patent 15 describes the use of these methods to produce hybrid REs for genome editing as well as other proteins that bind specific target DNA sequences for other applications. Patents 12, 14 and 15 are thus labeled as a cluster (C in the figure) which we refer to as Hybrid REs.

Another key step forward was the elucidation of the structure of zinc finger transcription factors revealing their modular zinc finger (ZF) structures responsible for DNA sequence specificity. This led to the idea that ZFs could be fused to a nuclease to create a hybrid RE with a novel DNA sequence specificity (36, 38, 39). In the late 90’s and early 00’s, patents 10 and 16 from Sangamo Biosciences describe the foundational invention of hybrid REs that fuse zinc finger DNA-binding domains with the FokI nuclease domain to create a zinc-finger nuclease (ZFN) capable of regulating or inactivating a target gene in its normal chromosomal context. These two patents and patent 17 constitute the ZFN labeled cluster D in figure 6.

The later discovery of transcription activator-like effectors (TALE) bacterial proteins that could, like zinc fingers, be engineered to create novel DNA binding specificities led to an analogous approach of fusing TALE binding domains to nucleases (36,38,39). Patents 21 and 26 from the Bonas group at Halle-Wittenberg University and Sangamo Biosciences respectively fused TALE domains to FokI nuclease to create TALE nucleases (TALENs) for genome engineering. More recent improvements in TALEN technologies by Sangamo are described in patents 22 and Patents 21, 22, 23 and 26 are thereby designated cluster F-TALENs.

In the late 1990’s, the discovery that the FokI nuclease is comprised of two monomers that require dimerization for nuclease activity led to the invention (Patent 17) of ZFN pairs comprising two monomers, each with a FokI half-cleavage domain and a zinc finger domain. ZFN pairs provided greater DNA target specificity because they require correct binding of two separate zinc fingers to reconstitute the nuclease activity of the FokI dimer (36).

In the past decade, patents 18, 19 and 20 describe the application of ZFN and TALEN genome engineering technologies for specific therapeutic purposes, such as to modulate PD1 gene expression for cancer immunotherapy (patent 18) or severe combined immunodeficiency (SCID) related genes (patents 19 and 20). Patents 24, 25, 27 and 28 from Factor Bioscience all describe extending the therapeutic application of ZFN, TALENs or CRISPR by therapeutic delivery of a synthetic RNA encoding the genome editing enzymes rather than DNA. In this way, the therapeutic nucleic acid is not incorporated into the genome potentially reducing the risk of unwarranted mutagenesis and limiting the therapeutic exposure to the lifespan of the RNA molecule.

The 20 patents just discussed and particularly the 12 (see Figure 6) that the technique identified as high persistence patents are clearly important patents as identified by other observers. The main path technique indicates that they are the most important in the overall development of genome editing prior to CRISPR. Therefore, we regard this small set of patents as the core technology preceding the CRISPR breakthrough but we do not regard all the rest of the 1373 patents in the set as unimportant since it is highly likely there are other quite important patents in the set.

#### CRISPR Roots Patents

The CRISPR roots patent set is different from the genome engineering patent set as it does not focus on a specific technical area (genome engineering) but instead backwardly traces *all* patented knowledge sources that have contributed to the emergence of CRISPR technology. Recall that the genome engineering patent set was carefully limited to chosen patent classes whereas the CRISPR roots set was subject to no such constraint. Additionally, all citations outside this selected set were ignored for the genome engineering main path analysis whereas the CRISPR roots includes *all* citations from the initial set of patents. The well-known and important phenomenon known as spillover means that the roots patent set will reflect broad sources of knowledge not included in the genome engineering domain.

The difference in breadth between the CRISPR roots and the genome engineering patent set is visible in the main path derived from the roots patent set. Figure 7 shows the result from application of the main path method to that patent set. Since this patent set is obtained starting with the citations by the currently published CRISPR patents, this knowledge network is constrained to end on the right at the CRISPR patents and the main path identifies patents that were particularly important in citations cascading back from these patents. The reasoning to develop this non-usual main path was simply to reduce the 1300+ patent set to the 50 most important ones so that it was possible to read and sort the patents.

**Figure 7.**
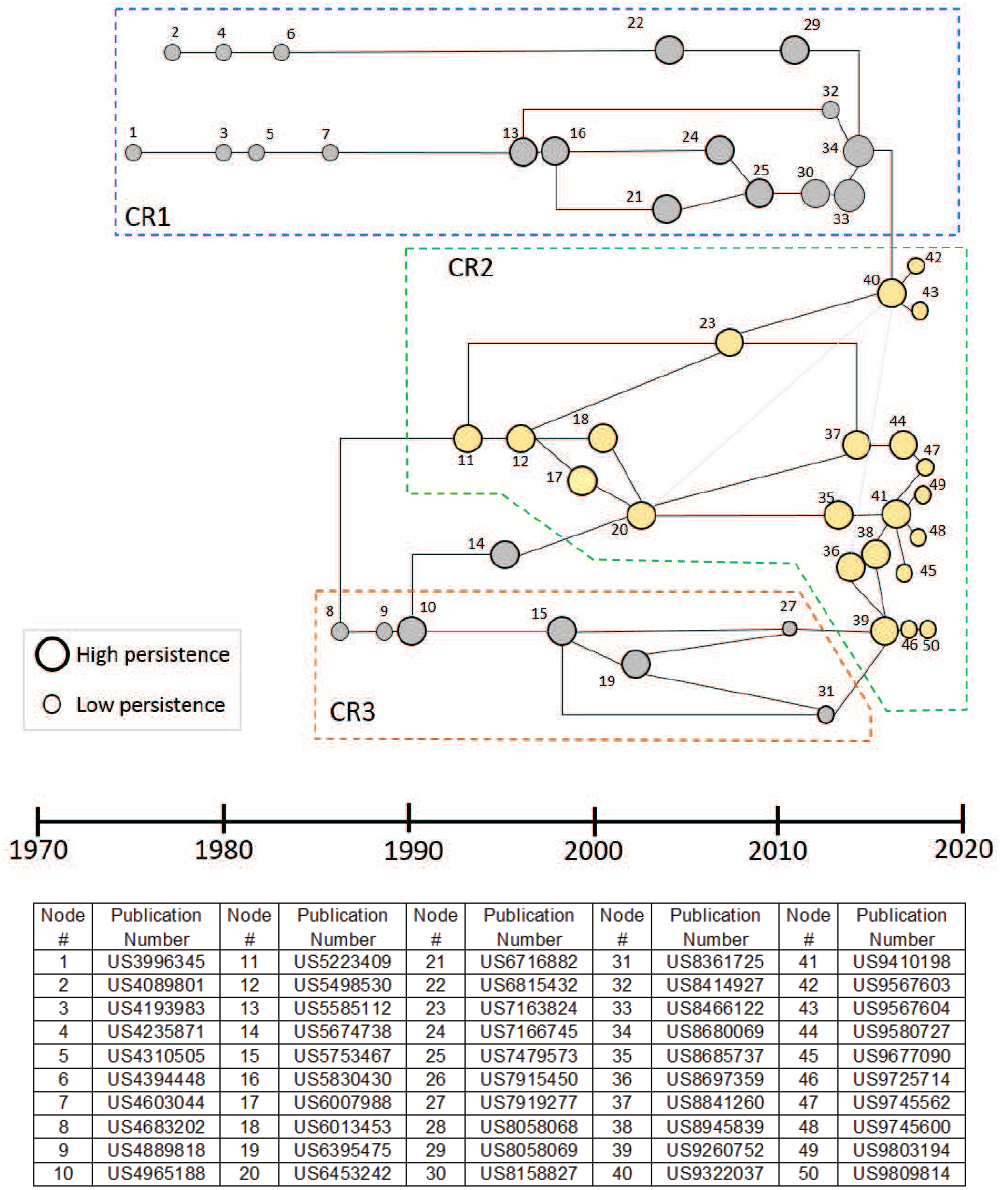
Main Path for the CRISPR roots showing patents on this knowledge trajectory from the CRISPR patents (gen 0), the patents cited by the CRISPR patents (gen 1) and the patents cited by gen 1 patents but not by CRISPR patents (gen 2). Three main paths (CR1, CR2, and CR3) have been identified. CR1: Technologies for introducing nucleic acid into mammalian cells; CR2: Genome engineering (including protein binding domains, ZFN and CRISPR); CR3: DNA finger printing and PCR. Labeled nodes represent patents shown in the table below the main path diagram. The node numbers increase along the time axis.

Like Figure 5, the main path network in Figure 7 also can be interpreted as consisting of three knowledge trajectories. At the top of the diagram is a large sequence of patents (CR1) that are concerned with delivery or the introduction of nucleic acid to mammalian cells. In the bottom part of Figure 7 are a set of patents (CR3) that involve DNA fingerprinting and demonstrate the pervasive impact of PCR on biotechnology as it emerges in the CRISPR context. The central main path or knowledge trajectory is genome engineering (CR2) which is connected to CR3 in 3 places and to CR1 in the link between patents 34 and 40. The presence of CR1 and CR3 paths in the roots main path demonstrates the broader scope of the CRISPR roots compared with the genome editing patent set. The patents in these paths were not in the genome engineering set by design but are shown in Figure 7 to play a prominent “spillover” role in the emergence of CRISPR.

Table 1 shows the ten patents with the highest normalized centrality (maximum = 1) from the CRISPR roots. Demonstrating the relative breadth in the CRISPR nucleus compared to the genome engineering patent set is the fact that *none* of these patents are in the genome engineering set. Instead, they include very important patents from the osmotic device domain, the ultrasound apparatus domain, nucleic acid methodology, crystal protein technology, and the drug delivery domain. With a minimum normalized centrality of > 0.986, these patents are highly important in their own domain and likely represent indirect or spillover technology essential to the development of CRISPR but are not on the genome engineering main path. Indeed, the second ranked patent in Table 1 is the very important/central PCR patent by Kary Mullis. It is probable that without PCR, there would be no CRISPR but this does not signify that this patent is on the main knowledge accumulation path leading to CRISPR. This result is similar to the broad scientific input that enabled CRISPR identified by Lander (9), by Doudna and Sternberg (10) and by Urnov (11) but the patents in Table 1 represent technological breadth not usually identified.

**Table 1:** The ten top-ranked patents from the CRISPR nucleus according to information centrality.

| Publication Number | Title | Publication Year | Standardized Inventor | Standardized Assignee | Centrality |
| --- | --- | --- | --- | --- | --- |
| US4210139 | Osmotic device with compartment for governing concentration of agent dispensed from device | 1980 | Higuchi Takeru | Alza Corp | 0.999883 |
| US4683202 | Process for amplifying nucleic acid sequences | 1987 | Mullis Kary B | Cetus Corp | 0.998493 |
| US4327725 | Osmotic device with hydrogel driving member | 1982 | Cortese Richard \| Theeuwes Felix | Alza Corp | 0.996192 |
| US4620546 | Ultrasound hyperthermia apparatus | 1986 | Aida Satoshi \| Matsumoto Kenzo \| Itoh Ayao \| ... | Toshiba | 0.993626 |
| US4959217 | Delayed/sustained release of macromolecules | 1990 | Sanders Lynda M \| Domb Abraham | Syntex Corp | 0.992252 |
| US5270163 | Methods for identifying nucleic acid ligands | 1993 | Gold Larry \| Tuerk Craig | Univ Research Corp, Boulder, CO | 0.988969 |
| US4900540 | Liposomes containing gas for ultrasound detection | 1990 | Ryan Patrick J \| Davis Michael A \| Melchior Donald L | Univ Of Massachusetts | 0.987955 |
| US5380831 | Synthetic insecticidal crystal protein gene | 1995 | Adang Michael J \| Rocheleau Thomas A \| Merlo Donald J | Mycogen Plant Science, Inc | 0.987837 |
| US4448885 | Bacillus thuringiensis crystal protein in Escherichia coli | 1984 | Schnepf H Ernest \| Whiteley Helen R | University Of Washington | 0.987605 |
| US5078994 | Microgel drug delivery system | 1992 | Nair Mridula \| Tan Julia S | Eastman Kodak | 0.986634 |

Although, as just emphasized, there are differences in the collection techniques and therefore in the results shown in Figures 5/6 and 7, there are also important similarities since both reflect the genome engineering work that preceded CRISPR. In this regard, we note that 5 of the top institutional owners of patents in the genome editing set are also in the top institutional owners of patents in the CRISPR roots set (compare Figure 2B and Figure 3B). Moreover, Table 2 shows 8 key patents in the main path of the genome engineering set that are also in CRISPR roots set. All 8 patents listed in Table 2 that are found in the CRISPR nucleus are also found in the GE2 (core genome editing) knowledge trajectory from the main path analysis of that domain. The node numbers in Table 2 are the ones given to these patents in Figure 6 which shows GE2 details and clusters. These 8 patents all relate to the foundational inventions of genome engineering prior to the discovery of CRISPR. As described above, patents 9 and 11 are inventions based on the discovery that the FokI restriction endonuclease is made of two separable DNA binding and cleavage domains. Patents 12, 14 and 15 describe methodological improvements in producing hybrid REs, while 10, 16 and 17 are related to the development of ZFNs as the first generally applicable hybrid REs for gene editing. The overlap between the patent sets is further illustration of the importance of earlier genome engineering technology to the development of CRISPR genome engineering despite the independent discovery of the original bacterial CRISPR viral resistance mechanism and all the very important but more distant knowledge represented in Table 1.

**Table 2:**
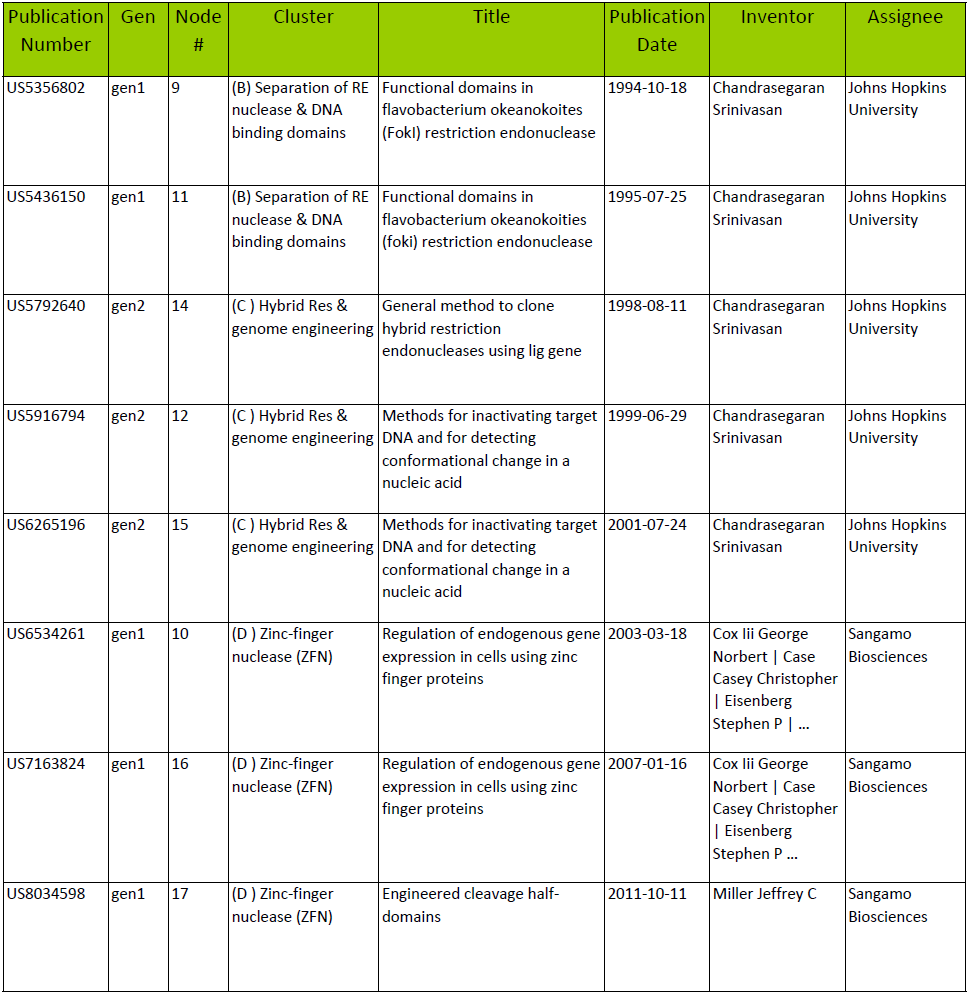
Eight key patents in the main path and core of genome editing which are also in the CRISPR roots set.

#### Performance Improvement results

Table 3 gives the results obtained when applying the k estimation algorithm described in the methods section (k is directly determined from the average centrality of the patent set) to the two patent sets. The first result is that the patent sets give different estimates of k (approximately x3 difference). Perhaps more significantly, both estimates are relatively low. We now briefly consider these two findings.

**Table 3:** The estimated annual improvement (k) in percentage for the genome engineering patent set (domain) and the CRISPR roots set.

| Patent Set (1970/01/01 – 2018/01/15) | k (Annual Improvement Rate %) |
| --- | --- |
| Genome engineering patent set | 3.6% |
| CRISPR Roots patent set | 9.5% |

Prior analysis of uncertainty in the k estimates (33, 34) indicates that +/- 50% uncertainty is a reasonable quantification for k +/- σ. This uncertainty in the estimate is consistent with empirical measurement of k (28, 29). Thus, the x3 difference in the estimated k values is probably not only due to uncertain estimates. Since these two sets of patents have large differences in what is included, significant differences in k are not unreasonable and could arise in various ways. One factor that appears likely to explain a large part of the differential result is the significantly larger breadth of the patents in the CRISPR roots which was discussed in the preceding section as reflecting the “spillover” patents in the roots that are not in the genome editing patent set. Such patents were not included in the domains where the empirical correlation was established (31,33) and would tend to distort k estimates for domains upwardly since patents cited from “farther afield” tend to be patents that are important in carrying information-that is have important new knowledge at their core- and thus have higher centrality than average. Since the genome engineering patent set has considerably lower average centrality (0.27) than the entire US patent set (0.5), including such patents in the set (as the roots set does) raises the overall k estimate. For example, the patents in Table 1 are the highest centrality patents from the roots set and were already seen as demonstrating breadth in the roots patent set.

Our second finding is that even the k value for the roots set is not very high in terms of what we now know about k values in various domains. Indeed, the average centrality of the genome engineering set is well below average (0.27) for USPTO patents and the average centrality of the CRISPR roots is higher (.43) but still below average for the entire US patent set which is equal to 0.5 (33).

## Discussion and Conclusions

Our first research objective was to determine what the patent record suggests relative to the relationship of CRISPR to prior technology-particularly pre-existing genome engineering. The results presented here (particularly Figure 6 and Table 3) show clearly that pre-existing genome engineering technology was essential to the emergence of CRISPR. There is close alignment of the qualitative history and the objective knowledge trajectory determination for the genome engineering patent set as shown by qualitatively known important patents being on the main path. Such agreement is what one would expect if the main path methodology and the patent selection methodology work as has been claimed previously (11, 12, 16, 17, 18). The present results thus offer some additional support to these prior claims.

The results in this paper go beyond confirming the expected importance of key earlier genome engineering developments on the emergence of CRISPR by demonstrating the quite broad array of technologies found in the CRISPR roots set (Table 1 and figure 7). The technologies playing an important and possibly essential role in CRISPR emergence include knowledge about PCR, knowledge from the osmotic device domain, from the ultrasound apparatus domain, from the crystal protein technology domain, and the drug delivery domain among many others. Such breadth is not unexpected from the prior knowledge of spillover but the specifics of the breadth is not usually determined. We note that qualitative histories tend to focus on the most direct technological path (or just the science) and thus do not begin to point to the technological breadth that may be essential to the emergence of highly novel and important technologies like CRISPR.

The results obtained in pursuit of our second objective (estimation of the rate of improvement for CRISPR) go well beyond anything done elsewhere. The estimate of the rate of technological performance improvement for CRISPR has been reported here and is the only estimate for *any* emerging technological domain to our knowledge. Since it is a first estimate of its kind, we must be careful to not over-claim significance and thus the following discussion should be considered preliminary until further patents emerge over time in the CRISPR domain and more importantly until other newly emerging technologies are studied by the techniques pioneered here. Although there has been some work on some emerging (but poorly defined) domains such as nanotechnology, this has not used the methods (main paths, roots investigation, rate estimation) applied herein to CRISPR. Most importantly, such domains typically have patents dating from many years back whereas the first CRISPR patent was in 2012. Studies of other emerging domains that we envision would concentrate on the initial 5-10 years after the initial patent.

Regarding the relatively low rate of performance improvement estimated for CRISPR, there are two topics worthy of such an early discussion. One is the potential importance of this observation to the evolving CRISPR story and another one is possible specific kind of performance improvement that is being estimated. As an initial remark on the significance of the observation in the evolving CRISPR story we do not believe low performance improvement rates mean that CRISPR is less important than it has been declared to be (1-10). However, we find it probable that the performance improvement being estimated is important rather than something to be ignored. One speculation is that the rate of improvement may relate to an unimportant metric; however, logical analysis of known results make this appear unlikely. It is unlikely first because it is usual (28) that *most intensive improvement rates in a domain are the same* within the normal variation so important and less important metrics tend to improve at the same rate. Moreover, some logical metrics for such a domain are clearly important; for example, a metric such as the increase in benefit (for example quality life years in a case like CRISPR) divided by the constraint (for example cost) is a likely relevant intensive metric that is improving at 3.5% (or possibly 9%) per year. To improve such a metric as Qualy/$ for CRISPR therapies from a very low starting point today will take solving multiple problems of harmful side effects while improving the ease with which genome engineering can be applied to a variety of human diseases. Thus (remembering our caveat about conclusions being preliminary), it is likely that important CRISPR based therapies will be appearing over many decades –not just in the next few years and that important developments in genome engineering will continue to build on and beyond CRISPR.

Our last conclusion from the research reported here is that the techniques used in the paper (main paths, comparing roots and the specific technological domain, k estimation) allow one to further understand specific technological developments very early after their emergence. However, we would like to stress that such objective methods are not a replacement for deep qualitative studies such as those by Lander, Doudna/Sternberg and Urnov (9,10,11) but instead are a valuable supplement. The supplement in this case is the clear *technological* breadth of CRISPR, the core gene editing patents linked to CRISPR, and the indication –even though preliminary- of relatively slow performance improvement of CRISPR.

## Acknowledgements

The authors gratefully acknowledge research support from Merck KGaA, Darmstadt, Germany and the help of Erik Vogan of MIT’s Industrial Liaison Office in initiating this research effort. The observations, description and opinions expressed here are those of the authors alone. We were in no way influenced by the granting organization or any other institution.

We also are grateful to Professor Giorgio Triulzi of the University of Los Andes (Columbia) for the centrality values for patents in both patent sets, to Professor Hyunseok Park of Hanyang university for the persistence main path software, and to Ritu Raj Lamsai of Deerwalk Institute for valuable inputs on patent searches and reading of patents.

